# CHMP2B regulates TDP-43 phosphorylation and proteotoxicity via modulating CK1 turnover independent of the autophagy-lysosomal pathway

**DOI:** 10.1101/2020.06.04.133546

**Authors:** Xing Sun, Xue Deng, Rirong Hu, Yongjia Duan, Kai Zhang, Jihong Cui, Jiangxia Ni, Qiangqiang Wang, Yelin Chen, Ang Li, Yanshan Fang

## Abstract

Protein inclusions containing phosphorylated TDP-43 are a shared pathology in several neurodegenerative diseases including amyotrophic lateral sclerosis (ALS) and frontotemporal dementia (FTD). However, most ALS/FTD patients do not have a mutation in TDP-43 or the enzymes directly regulating its phosphorylation. It is intriguing how TDP-43 becomes hyperphosphorylated in each disease condition. In a genetic screen for novel TDP-43 modifiers, we found that knockdown (KD) of *CHMP2B*, a key component of the endosomal sorting complex required for transport (ESCRT) machinery, suppressed TDP-43-mediated neurodegeneration in *Drosophila*. Further investigation using mammalian cells indicated that *CHMP2B* KD decreased whereas its overexpression (OE) increased TDP-43 phosphorylation levels. Moreover, a known FTD-causing mutation *CHMP2B*^*intron5*^ promoted hyperphosphorylation, insolubility and cytoplasmic accumulation of TDP-43. Interestingly, CHMP2B did not manifest these effects by its well-known function in the autophagy-lysosomal pathway. Instead, the kinase CK1 tightly regulated TDP-43 phosphorylation level in cells, and CHMP2B OE or CHMP2B^Intron5^ significantly decreased ubiquitination and the turnover of CK1 via the ubiquitin-proteasome (UPS) pathway. Finally, we showed that CHMP2B protein levels increased in the cerebral cortices of aged mice, which might underlie the age-associated TDP-43 pathology and disease onset. Together, our findings reveal a molecular link between the two ALS/FTD-pathogenic proteins CHMP2B and TDP-43, and provide an autophagy-independent mechanism for CHMP2B in pathogenesis.

**SIGNIFICANCE STATEMENT:** TDP-43 and CHMP2B are both ALS/FTD-associated proteins. Protein aggregations containing phosphorylated TDP-43 are a pathological hallmark of ALS/FTD; however, it is unclear how increased phosphorylation of TDP-43 occurs in diseases. The pathogenesis of CHMP2B has mainly been considered as a consequence of autophagy-lysosomal dysfunction. Here, we reveal that increase of CHMP2B levels (which occurs in aged mouse brains) or expression of the disease-causing mutation CHMP2B^Intron5^ promotes TDP-43 hyperphosphorylation, insolubility and cytoplasmic mislocalization. This effect is independent of the autophagy-lysosomal pathway but rather relies on the proteasome-mediated turnover of the kinase CK1 that phosphorylates TDP-43. Together, we provide a new molecular mechanism of CHMP2B pathogenesis by linking it to TDP-43 pathology via CK1.

## INTRODUCTION

TAR DNA-binding protein 43 (TDP-43) is a nuclear RNA/DNA-binding protein that can shuttle between the nucleus and the cytoplasm. Under normal conditions, TDP-43 participates in the assembly and function of various ribonucleoprotein (RNP) complexes and plays an important role in regulating RNA processing and metabolism (Chen-Plotkin et al., 2010; Cohen et al., 2011; Lee et al., 2012). In disease, abnormal TDP-43 protein inclusions are found in ∼97% ALS and ∼45% FTD patients (Ling et al., 2013; Tan et al., 2017). In fact, TDP-43 is a major pathological protein linked to a spectrum of neurological disorders, collectively known as TDP-43 proteinopathies, which include not only ALS and FTD but also Alzheimer’s disease (AD), dementia with Lewy bodies and polyglutamine diseases (Neumann et al., 2007; Higashi et al., 2007; Arai et al., 2009; Toyoshima et al., 2014; Chang et al., 2016). TDP-43 pathology is characterized by hyperphosphorylation of the protein at serine 409 and 410 (pS409/410) (Hasegawa et al., 2008; Neumann et al., 2009). Despite being a shared pathology, mutations in *TARDBP* (the gene encoding TDP-43) only account for 1∼5% family ALS and about 1% family FTD (Ji et al., 2017), and none of the known ALS/FTD causal genes encodes a kinase or phosphatase that directly regulates TDP-43 phosphorylation (Pottier et al., 2016; Corcia et al., 2017). As such, it is largely unknown how the pathological processes that promote TDP-43 phosphorylation and aggregation occur in most ALS/FTD cases.

Phosphorylation of TDP-43 is closely related to its insolubility, aggregation tendency, cytoplasmic mislocalization and the pathogenesis. For example, phosphorylated TDP-43 (pTDP-43) is more detergent-insoluble, has a longer half-life, forms high-molecular weight oligomers and fibrils, and accumulates cytoplasmic aggregations in cells (Liachko et al., 2010; Zhang et al., 2010; Choksi et al., 2014; Nonaka et al., 2016; Yamashita et al., 2016), and plasma pTDP-43 levels correlate with the extent of brain pathology in FTD patients (Foulds et al., 2009). Casein kinase 1 (CK1) is a family of serine/threonine-selective kinases, which phosphorylate key proteins and function in developmental signaling such as Hedgehog and Wnt pathways as well as in regulating circadian rhythms and cellular metabolism (Cheong and Virshup, 2011; Cruiciat, 2014; Jiang, 2017). Recently, mounting evidence suggests the relevance of CK1 in human diseases such as cancer and neurodegenerative disorders (Perez et al., 2011; Cozza and Pinna, 2016). In particular, CK1 could phosphorylate TDP-43 *in vitro* (Hasegawa et al., 2008; Kametani et al., 2009) and the phosphorylated epitopes generated by CK1 could be strongly recognized by TDP-43 antibodies in immunohistological examinations of FTD and ALS brains (Hasegawa et al., 2008). In cell and animal models, expression of a hyperactive form of CK1δ promoted TDP-43 phosphorylation (Nonaka et al., 2016), whereas CK1 inhibitors were shown to reduce TDP-43 phosphorylation and suppress its toxicity in mammalian cell cultures, fly and mouse neurons, and human cells derived from FTD and ALS patients (Salado et al., 2014; Alquezar et al., 2016; Martinez-Gonzalez et al., 2020). Thus, modulating CK1 activity may be a potential therapeutic approach for treating FTD and ALS. However, whether and how CK1 is involved in the disease pathogenesis is unclear.

*Charged multivesicular body protein 2B* (*CHMP2B*) is another gene whose mutations are associated with both ALS and FTD (Skibinski et al., 2005; Parkinson et al., 2006; Cox et al., 2010). It encodes the ESCRT-III subunit protein CHMP2B, which plays a vital role in endolysosomal trafficking, vesicle fusion and autophagic degradations (Urwin et al., 2009; Henne et al., 2011). Among the disease-associated mutations in the *CHMP2B* gene, the most studied is CHMP2B^Intron5^ in FTD linked to chromosome 3 (FTD-3). It is a single nucleotide (G→C) mutation in exon 6, causing aberrant splicing inclusive of the 201 bp of intron 5, which contains a stop codon and leads to a truncation of the C-terminus of CHMP2B. The resulting CHMP2B^Intron5^ protein lacks the important binding site for Vps4, whose recruitment is required to initiate membrane abscission and the disassembly of the ESCRT-III complex. Besides, as the CHMP2B^Intron5^ protein loses 36 amino acids from the acidic C-terminus, its self-binding to the basic N-terminus is significantly reduced, which disrupts the normal autoinhibition in CHMP2B (Urwin et al., 2009; Isaacs et al., 2011; Krasniak and Ahmad, 2016). Dysfunction of the ESCRT-III complex leads to accumulation of endosomes and autophagosomes, which has been evident in a variety of cell and animal models (Lee et al., 2007; van der Zee et al., 2008; Urwin et al., 2010; Ghazi-Noori et al., 2012; Clayton et al., 2015). Thus, the CHMP2B pathogenesis has mainly been considered as a downstream consequence of the disrupted autophagic and endolysosomal pathway. In addition, TDP-43-immunoreactive inclusion bodies were found in motor neurons and glia of ALS patients containing *CHMP2B* mutations (ALS17) (Cox et al., 2010). However, it remains unclear whether CHMP2B and TDP-43 are molecularly linked and how this contributes to ALS/FTD pathogenesis.

In this study, we identify CHMP2B as a novel modifier of TDP-43 neurotoxicity in a fly genetic screen. Further investigation reveals a striking role of CHMP2B in regulating TDP-43 phosphorylation levels, protein solubility and subcellular distribution. Unexpectedly, although manipulation of CHMP2B levels or expression of CHMP2B^intron5^ indeed impacts on the autophagy-lysosomal proteolysis, disruption of this pathway does not alter TDP-43 phosphorylation levels, suggesting an autophagy-independent mechanism. Rather, it is the kinase CK1 that mediates the modifying effect of CHMP2B on TDP-43 phosphorylation and cytotoxicity, as CHMP2B modulates the abundance of CK1 via the UPS-dependent protein turnover. Together, we propose that a molecular axis of “CHMP2B–CK1–TDP-43” may be involved in the pathogenesis of ALS/FTD and related diseases.

## RESULTS

### Downregulation of *CHMP2B* alleviates TDP-43 neurotoxicity in a *Drosophila* model

To identify unknown players involved in TDP-43-mediated neurodegeneration, we conducted a genetic screen for new modifiers of TDP-43 neurotoxicity using a *Drosophila* model that expressed human TDP-43 (*hTDP-43*) with the binary Gal4-UAS system. Transgenic expression of *hTDP-43* in the fly eyes (by a GMR-Gal4 driver) caused age-dependent degeneration, evident by rough surface, loss of pigment cells and swelling of the compound fly eyes (Figure 1A-D; Figure S1A-B).

**Figure 1.**
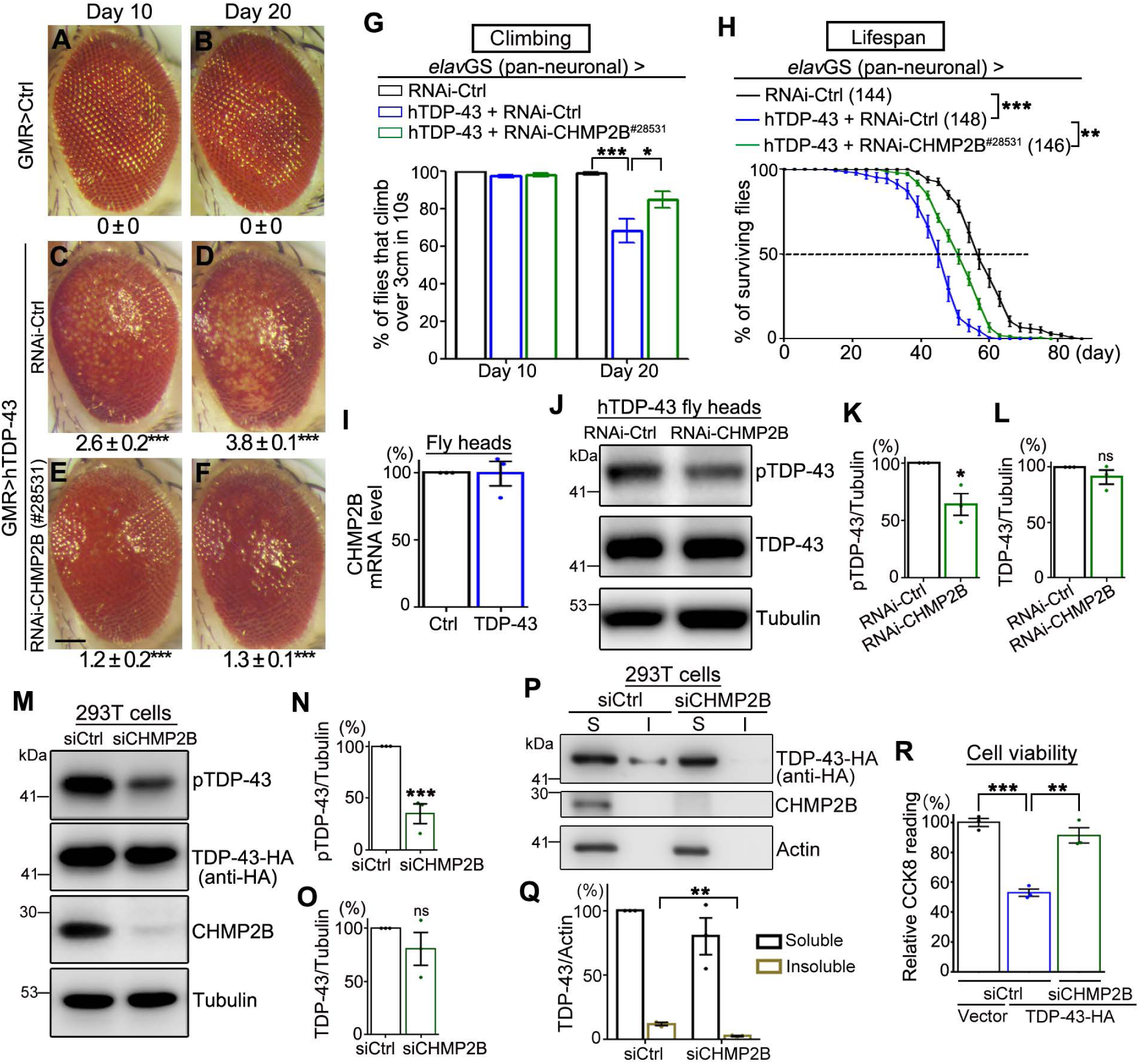
Identification of RNAi-*CHMP2B* as a suppressor of TDP-43-mediated cytotoxicity in *Drosophila* and mammalian cells. (**A-F**) Representative z-stack images of the fly eyes expressing the indicated transgenes (by GMR-Gal4) at Day 10 or Day 20. The average degeneration score (mean ± SEM) and the statistical significance compared to the UAS-*lacZ* control (Ctrl) line are indicated at the bottom of each group. (**G**) The climbing capability of the flies expressing the indicated transgenes in adult neurons (by *elav*GS) is evaluated as the average percentage of flies that climb over 3 cm in 10 seconds. (**H**) Lifespan assays of the flies with adult-onset, neuronal expression (by *elav*GS) of the transgenes as indicated. The numbers of flies tested in each group are indicated. The RNAi-*luciferase* fly line is used as a control (RNAi-Ctrl). (**I**) qPCR analysis of the mRNA levels of *CHMP2B* in the heads of TDP-43 flies. The mRNA levels are normalized to *actin* and shown as average percentage to that of the UAS-*lacZ* (Ctrl) group. (**J-L**) Representative images (J) and quantifications (K-L) of the Western blot analysis of the levels of S409/410 pTDP-43 (K) or total TDP-43 protein (L) in the fly heads. The protein levels are normalized to Tubulin and shown as percentage to the control group. (**M-O**) Representative images (M) and quantifications (N-O) of the Western blot analysis of pTDP-43 levels (N) or total TDP-43-HA protein levels (O) in 293T cells. (**P-Q**) Representative western blot images (P) and quantification (Q) of TDP-43-HA proteins in the soluble (S, supernatants in RIPA) and insoluble fractions (I, pellets resuspended in 9 M of urea) of 293T cell lysates. All protein levels are normalized to Actin in the soluble fractions. (**R**) The viability of 293T cells transfected with the empty vector or TDP-43-HA together with scrambled siRNA (siCtrl) or siRNA against CHMP2B (siCHMP2B) is assessed using the CCK-8 assay. Mean ± SEM, n = ∼10 eyes each group in (A-F), ∼20 flies/vial and ∼10 vials/group in (G), and 3 in (I, K-L, N-O, Q-R). Statistical significance was determined by Student’s *t*-test (A-G, I, K-L, M-N, R) and two-way ANOVA (H) at \**p* < 0.05, \*\**p* < 0.01 and \*\*\**p*<0.001; ns, not significant. Scale bar: 100 μm.

In the screen, two transgenic fly lines (#28531 and #38375) showed suppression of *hTDP-43*-induced eye degeneration (Figure 1E-1F and Figure S1C-S1D), which turned out to be two independent RNAi lines of the fly gene *CHMP2B*. Further examination revealed that downregulation of *CHMP2B* with an inducible pan-neuronal *elav*-GeneSwitch (*elav*GS) driver markedly mitigated *hTDP-43*-induced climbing deficits during aging (Figure 1G; Figure S1E). Moreover, expression of *hTDP-43* in the adult fly neurons shortened the lifespan by 20.7% (median life, from 57.9 ± 1.2 d to 45.9 ± 1.1 d), while KD of *CHMP2B* increased the lifespan of the TDP-43 flies by 14.4% (median life, from 45.9 ± 1.1 d to 52.5 ± 0.8 d) (Figure 1H). The other RNAi-*CHMP2B* transgenic fly line also partially rescued the shortened lifespan of the TDP-43 flies though to a lesser extent (Figure S1F). Of note, KD of *CHMP2B* did not increase the climbing capability or lifespan of control flies (Figure S1G-H), indicating that downregulation of *CHMP2B* did not have a general beneficial effect; rather, it specifically modified TDP-43-mediated neurodegeneration in flies.

### KD of *CHMP2B* decreases TDP-43 phosphorylation levels and cytotoxicity in flies and mammalian cells

We then set out to probe for the molecular mechanism underlying the modifying effects of *CHMP2B* on TDP-43 neurotoxicity. Since TDP-43 plays a vital role in regulating RNA metabolism and protein homostasis (Cohen et al., 2011; Lee et al. 2012), it would be reasonable to hypothesize that TDP-43 might affect *CHMP2B* mRNA levels and thus correction of this alteration by RNAi-*CHMP2B* could rescue the phenotypes of TDP-43 flies. However, real-time quantitative PCR (qPCR) analysis of the fly heads did not show a significant alteration of the mRNA levels of *CHMP2B* (Figure 1I). Then, the alternative hypothesis was that CHMP2B might regulate TDP-43 instead. Considering the known function of CHMP2B and ESCRT in autophagy and protein turnover in cells, we examined the protein abundance of TDP-43 in the RNAi-*CHMP2B* flies by western blotting. Again, no significant change of TDP-43 levels was detected (Figure 1J, 1L). Unexpectedly, the phosphoyrlation of TDP-43 at S409/410 (pS409/410), a disease hallmark of TDP-43 pathology (Hasegawa et al., 2008; Neumann et al., 2009), was significantly decreased (Figure 1J-1K).

We then extended our study to mammlian cells. First, we confirmed that TDP-43 OE did not alter CHMP2B mRNA or protein levels in human 293T cells (Figure S2A-S2C). Next, to test whether CHMP2B modulated TDP-43 levels, we downregulated CHMP2B by small interference RNA (siRNA). Consistent with the fly data, KD of CHMP2B decreased pTDP-43 levels without significantly affecting TDP-43 protein abundance in mammlian cells (Figure 1M-O). siRNA of CHMP2B (siCHMP2B) also reduced the presense of TDP-43 in the detergent-insoluble protein fraction (Figure 1P-Q), which was consistent with the previous reports that pTDP-43 showed decreased solubility (Nonaka et al., 2016; Yamashita et al., 2016). More importantly, we examined the cell viability using a Cell Counting Kit-8 (CCK8) assay, which confirmed that KD of CHMP2B significantly suppressed TDP-43 OE-induced cytotoxicity in human cells (Figure 1R).

### OE of CHMP2B or CHMP2B^intron5^ promotes pathological alterations of TDP-43 biochemical properties

Next, we examined how upregulation of CHMP2B or expressoin of the disease-causing mutation CHMP2B^intron5^ impacted on TDP-43 in 293T cells. We showed that OE of CHMP2B increased TDP-43-mediated cytotoxicity, and co-transfection of CHMP2B^intron5^ drastically enhanced the loss of cell viability (Fig. 2A). Western blot analyses indicated that pTDP-43 levels of both endogenous (Figure 2B-2D) and transiently expressed TDP-43 (Figure 2E-2G) were substantially increased by co-expression of CHMP2B or CHMP2B^intron5^. Furthermore, although the protein levels of RIPA-soluble TDP-43 were largely unchanged, there was a dramatic increase of the insoluble fractions of TDP-43 protein when CHMP2B or CHMP2B^intron5^ was co-expressed (Figure 2H-2I).

**Figure 2.**
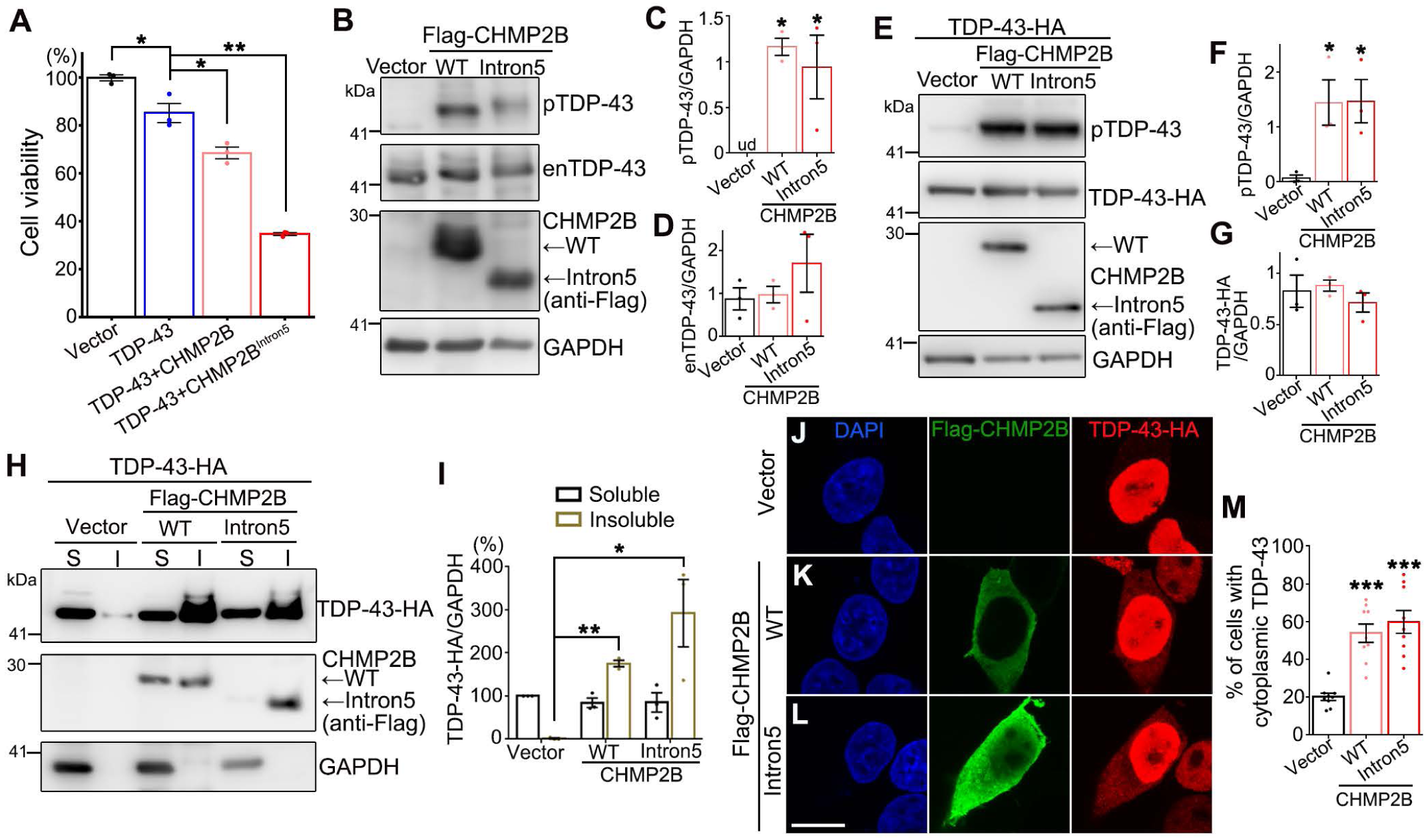
OE of CHMP2B or CHMP2B^Intron5^ promotes TDP-43 cytotoxicity, phosphorylation, insolubility and cytoplasmic localization. (**A**) Co-transfection of WT or Intron5 CHMP2B enhances TDP-43-induced reduction of cell viability in 293T cells. (**B-G**) Representative Western blot images (B, E) and quantifications (C-D, F-G) of the phosphorylation levels and protein abundance of endogenous TDP-43 (enTDP-43) (B-D) or transiently expressed TDP-43-HA (E-G) in 293T cells. ud, undetected. (**H-I**) Representative Western blot images (H) and quantifications (I) of TDP-43 protein in the soluble (S, supernatants in RIPA) and insoluble fractions (I, pellets resuspended in 9 M of urea) of 293T cell lysates. Cells transfected with the empty vector are used as a control. All protein levels are normalized to GAPDH in the soluble fractions. (**J-M**) Representative images (J-L) and quantification (M) of the immunocytochemistry analyses of TDP-43 subcellular distribution in 293T cells co-transfected with WT or Intron5 CHMP2B as indicated. Mean ± SEM, n = 3 replicates in (A, C-D, F-G, I) and ∼200 cells each group of pooled results from 3 independent repeats in (M). \**p* < 0.05, \*\**p* < 0.01, \*\*\**p*<0.001; one-way ANOVA; ud, undetectable. Scale bar: 10 μm.

Cytoplasmic mislocalization of TDP-43 is a common feature of TDP-43 pathology and is thought to be assoicated with its hyperphosphorylation (Lee et al., 2012; Nonaka et al., 2016). We then conducted immunocytochemistry analysis to examine whether CHMP2B affected the subcellular distribution of TDP-43. In normal cells, TDP-43 was predominantly nuclear (Figure 2J). Co-expression of CHMP2B (Figure 2K) or CHMP2B^intron5^ (Figure 2L) caused significant cytoplasmic translocation and/or accumulation of TDP-43, as more cells showed cytoplasmic TDP-43 (Figure 2M). Together, these results indicated that increase of CHMP2B levels in cells, by OE of either wildtype (WT) CHMP2B or CHMP2B^Intron5^, promoted the cytotoxicity as well as several key pathological alterations of TDP-43, including increased phosphorylation levels, insolubility and abnormal cytoplasmic localization.

### The autophagy-lysosomal pathway may not mediate the modifying effects of CHMP2B on TDP-43 phosphorylation

CHMP2B and the ESCRT complex play an important role in autophagy and endolysosomal pathways, and the CHMP2B^Intron5^ mutation was assoicated with accumulation of autophagsosomes in cells (Lee et al., 2007). We thus hypothesized that dysfunction of the autophagy-lysosomal pathway by CHMP2B mutations might account for the increased phosphorylation levels and the other biochemical alterations of TDP-43. To test the hypothesis, we examined the levels of autophagy markers including the microtubule-associated protein light chain 3 (LC3)-II and P62. LC3-II is formed by conjugating with phosphatidylethanolamine during autophagy when autophagosomes engulf cytoplasmic components (Kabeya et al., 2000); P62 is a substrate preferentially degraded by autophagy and thus its levels are often used as an indication of the function of autophagic-lysosomal proteolysis (Bjørkøy et al., 2005). We found that both KD and OE of CHMP2B increased the autophagy marker LC3-II protein levels (Fig. 3A-3B, 3D-3E), while KD of CHMP2B also increased another autophagy protein P62 (Fig. 3A, 3C, 3D, 3F), indicating that both down and up-regulation of CHMP2B led to autophagy-lysosomal dysfunction. Moreover, expression of the FTD3-assoicated CHMP2B^intron5^ mutation caused a drastic increase of both LC3-II and P62 proteins, which was consistent with the previous report that CHMP2B^Intron5^ mutation disrupted autophagic flux (Lee et al., 2007).

**Figure 3.**
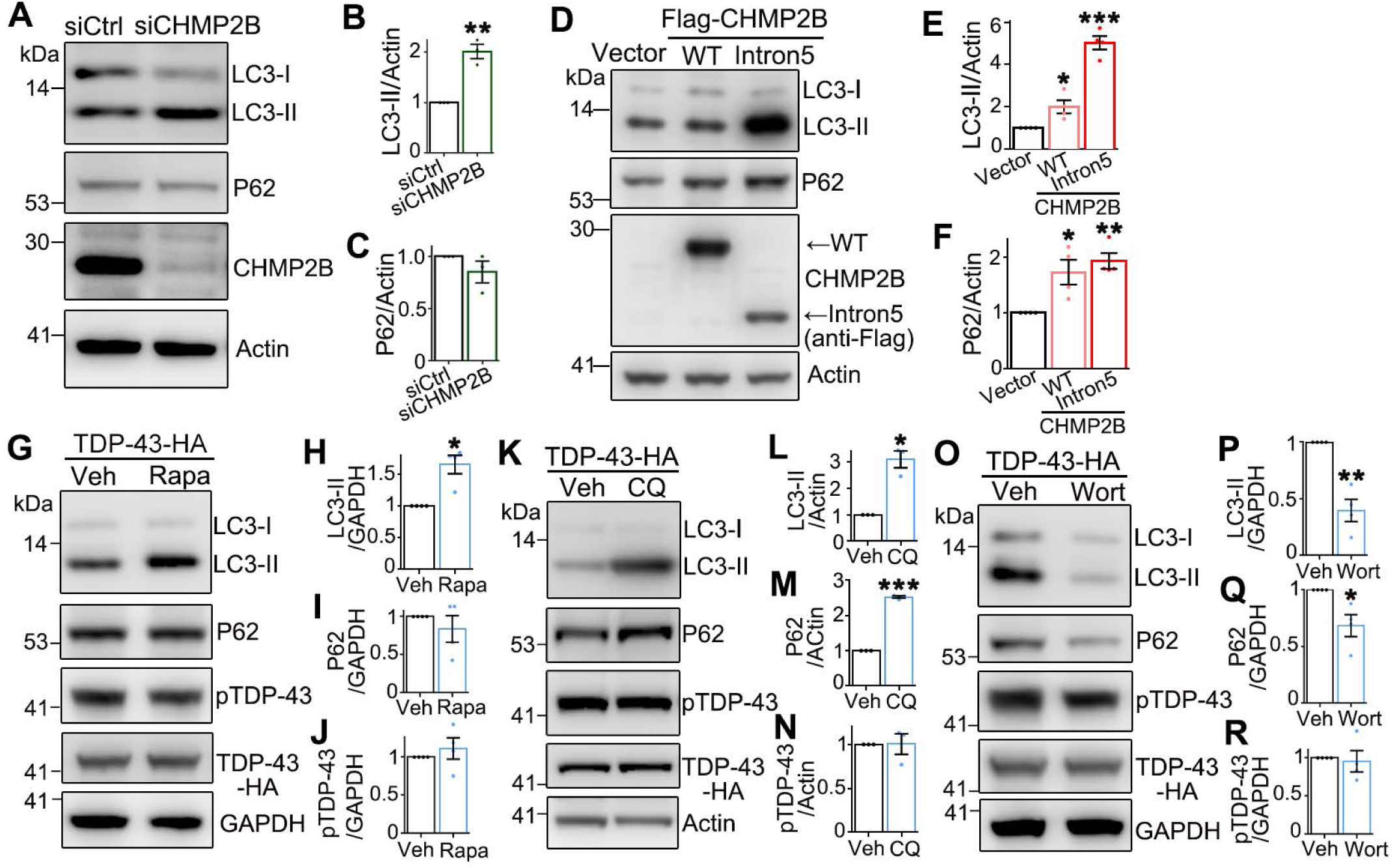
CHMP2B regulates autophagy but autophagic-lysosomal dysfunction does not affect pTDP-43 levels. (**A-F**) Representative images (A, D) and quantifications (B-C, E-F) of Western blot analyses of 293T cells treated with siCHMP2B (A-C) or OE of WT CHMP2B or CHMP2B^Intron5^ (D-F). Scrambled siRNA is used as a control (siCtrl). (**G-R**) Cells transiently expressing TDP-43-HA were treated with rapamycin (Rapa, 100 nM, 1 h) (G-J), chloroquine (CQ, 10 μM, 12 h) (K-N), or wortmannin (Wort, 50 nM, 1 h) and analyzed by Western blotting. Vehicle (Veh) controls: DMSO for Rapa and Wort, and PBS for CQ. Mean ± SEM, n = 3∼4 independent repeats in (B-C, E-F, H-J, L-N, P-R). \**p* < 0.05, \*\**p* < 0.01, \*\*\**p*<0.001; Student’s *t*-test except for (E-F, one-way ANOVA).

To test if autophagy dysfunction underlied the modifying effect of CHMP2B on TDP-43 phosphorylation states, we set out to determine how disturbance of the autophagy-lysosomal pathway impacted on pTDP-43 levels. To do this, we treated cells with several commonly used autophagy inducers and inhibitors, including rapamycin (a well studied mTOR inhibitor that activates autophagy), chloroquine (which inhibits both fusion of autophagosomes with lysosomes and lysosomal protein degradation), and wortmannin (an irreversible inhibitor of phosphatidylinositol 3-kinase (PI3K) whose activation is required for autophagy initiation) (Klionsky et al., 2016). We showed that activation of autophagy with rapamycin led to an increase of LC3-II but not P62 levels (Figure 3G-3I), which mimicked the effect of siCHMP2B (Figure 3A-3C); inhibition of the autophagy-lysosomal degradation by chloroquine led to the accumulation of both LC3-II and P62 (Figure 3K-3M), which mimicked the effect of OE of CHMP2B and CHMP2B^Intron5^ (Figure 3D-3F). In addition, the use of wortmannin potently inhibited autophagy, evident by a substantial decrease in both LC3-II and P62 levels (Figure 3O-3Q). Unexpectedly, however, in none of the above tests did we detect a significant change of pTDP-43 levels (Figure 3G-3R). Thus, it was unlikely that CHMP2B modulated TDP-43 phosphorylation states via the autophagy-lysosomal pathway.

### CK1 is involved in the regulation of TDP-43 phosphorylation and cytotoxicity by CHMP2B

To seek for the alternative mechanism that mediated the modifying effect of CHMP2B on TDP-43 phosphorylation, next we focused on the protein kinase CK1, which was previously demonstrated to phosphorylate TDP-43 *in vitro* (Hasegawa et al., 2008; Kametani et al., 2009) and its inhibitors may be of therapeutic potentials for treating FTD and ALS (Salado et al., 2014; Alquezar et al., 2016; Martinez-Gonzalez et al., 2020). In this study, we showed that CK1 such as CK1α and CK1δ could interact with TDP-43 in cells (Figure S3). And, OE of CK1α or CK1δ increased pTDP-43 levels (Figure 4A-4B), and promoted abnormal cytoplasmic localization of TDP-43 (Figure 4C-4D). More importantly, KD of CK1α and CK1δ by siRNA decreased pTDP-43 levels (Figure 4E-4F) and suppressed the cytotoxicity of TDP-43 (Figure 4G). Of note, due to the critical role of CK1 in regulating the functions and survival of cells, the amounts of transfection plasmids (10 ng of CK1α and 5 ng of CK1δ, Figure S4A) and siRNAs (0.8 nM siCK1α and 0.16 nM si CK1δ, Figure S4B) were carefully determined so that OE or KD of of CK1α and CK1δ did not manifest significant toxicity by themselves in the above and following cell viability assays in this study.

**Figure 4.**
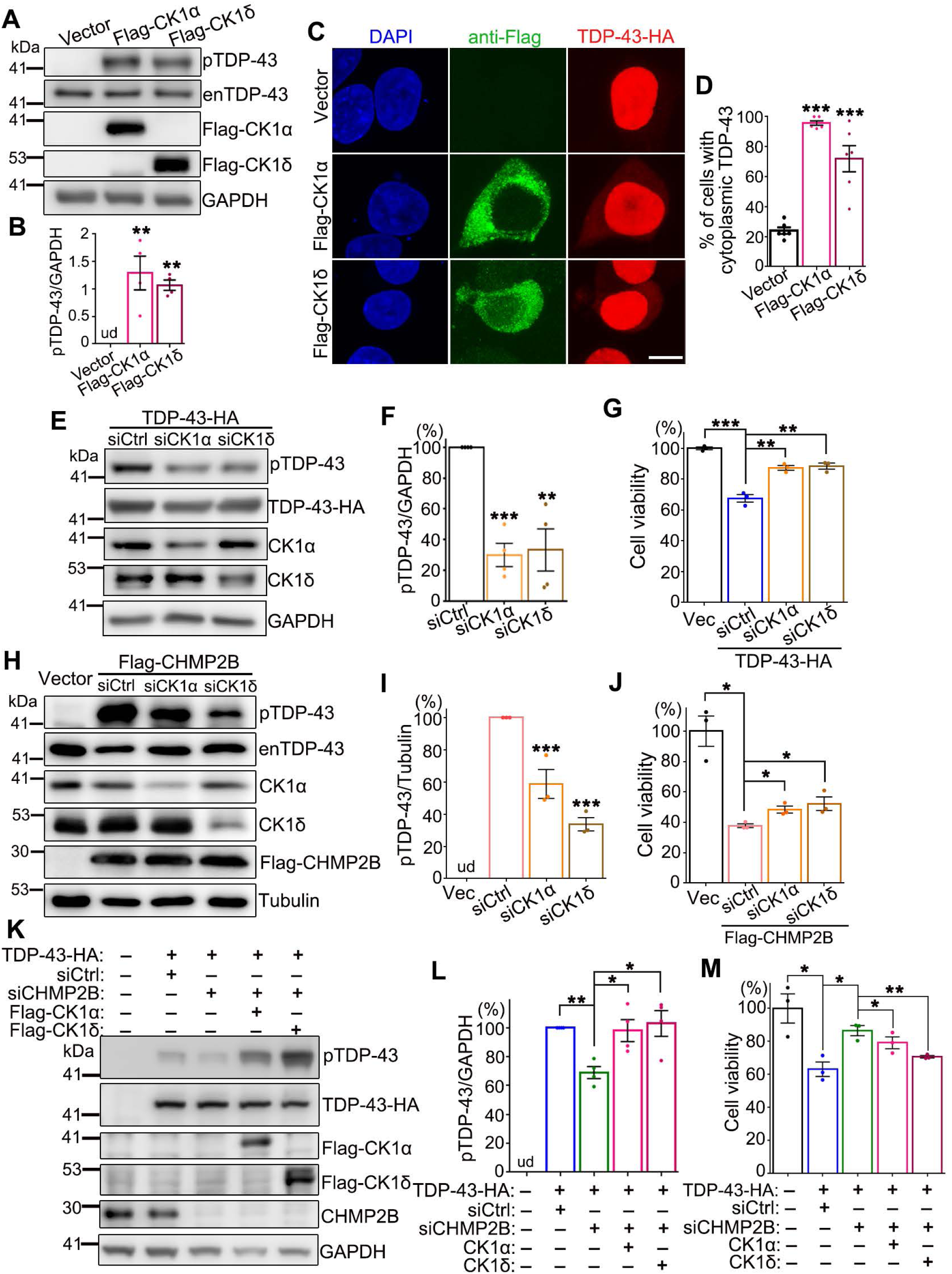
CK1 mediates CHMP2B-induced TDP-43 hyperphosphorylation and cytotoxicity. **(A-B)** Representative images (A) and quantifications (B) of Western blot analysis of the phosphorylation levels of endogenous TDP-43 (enTDP-43) in 293T cells transiently transfected of Flag-CK1α or Flag-CK1δ. **(C-D)** Representative confocal images (C) and quantifications (D) of subcellular localization of TDP-43-HA in cells co-transfected with Flag-CK1α or Flag-CK1δ. **(E-F)** Representative Western blot images (E) and quantifications (F) of the phosphorylation levels of TDP-43-HA in 293T cells treated with siCK1α or siCK1δ. **(G)** The cell viability assay indicates KD of CK1 suppresses the cytotoxicity of TDP-43. **(H-I)** CHMP2B OE-induced hyperphosphorylation of endogenous TDP-43 can be rescued by siCK1α or siCK1δ. (**J**) siCK1α or siCK1δ suppresses CHMP2B OE-induced cytotoxicity. **(K-L)** Reduction of pTDP-43 levels by siCHMP2B is abolished by OE of CK1α or CK1δ. **(M)** OE of CK1α or CK1δ significantly diminishes the mitigating effect of siCHMP2B on TDP-43-mediated cytotoxicity. Of note, the amount of the transfection plasmids and siRNAs of CK1α and CK1δ is used at the minimal sufficiency as indicated to avoid the cytotoxic of OE or KD of CK1 by itself (also see Figure S3). Cells transfected with the empty vector (Vec) or scrambled siRNA (siCtrl) are used as the controls in the above assays. Mean ± SEM, n = 3∼4 in (B, F, G, I, J, L, M) and ∼200 cells each group of pooled results from 3 independent repeats in (D). \**p* < 0.05, \*\**p* < 0.01, \*\*\**p*<0.001; one-way ANOVA; ud, undetectable. Scale bar: 10 μm.

Next, we examined whether manipulation of CK1 levels affected the modifying effects of CHMP2B on TDP-43 phosphorylation. Indeed, downregulation of CK1α or CK1δ not only suppressed CHMP2B-induced TDP-43 hyperphosphorylation (Figure 4H-4I) but also partially rescued the cytotoxicity caused by CHMP2B OE (Figure 4J). Consistentely, upregulation of CK1α or CK1δ abolished the mitigating effects of siCHMP2B on TDP-43 phosphorylation levels (Figure 4K-4L) and cytotoxicity (Figure 4M). Together, these data indicated that the modulation of TDP-43 phosphorylation and cytotoxicity by CHMP2B involved the function of CK1.

### CHMP2B regulates CK1 abundance via the UPS-mediated protein turnover

In the earlier experiments, we noticed that even mild OE or KD of CK1α or CK1δ was sufficient to alter pTDP-43 levels (10 ng of CK1α and 5 ng of CK1δ in Figure 4H-4I, and 0.8 nM siCK1α and 0.16 nM si CK1δ in Figure 4K-4L; also see Figure S4), suggesting that the protein levels of CK1 were critical in determining TDP-43 phosphorylation states in cells. This prompted us to test whether CHMP2B had a major regulatory effect on CK1 abundance. OE or KD of CHMP2B did not significantly change the mRNA levels of CK1α or CK1δ (Figure S5A-S5D). In contrast, the protein levels of CK1α and CK1δ were significantly reduced by KD of CHMP2B (∼50% reduction, Figure 5A-5C), whereas their protein levels were drastically increased by OE of CHMP2B (2∼3 folds increase, Figure 5D-5F). Next, we measured the protein turnover rate of CK1α and CK1δ by the pulse-chase assay with cycloheximide (CHX) to inhibit protein translation. We found that the turnover of CK1α and CK1δ was accelerated by siCHMP2B (Figure 5G-5I) but substantially impeded by OE of CHMP2B (Figure 5J-5L).

**Figure 5.**
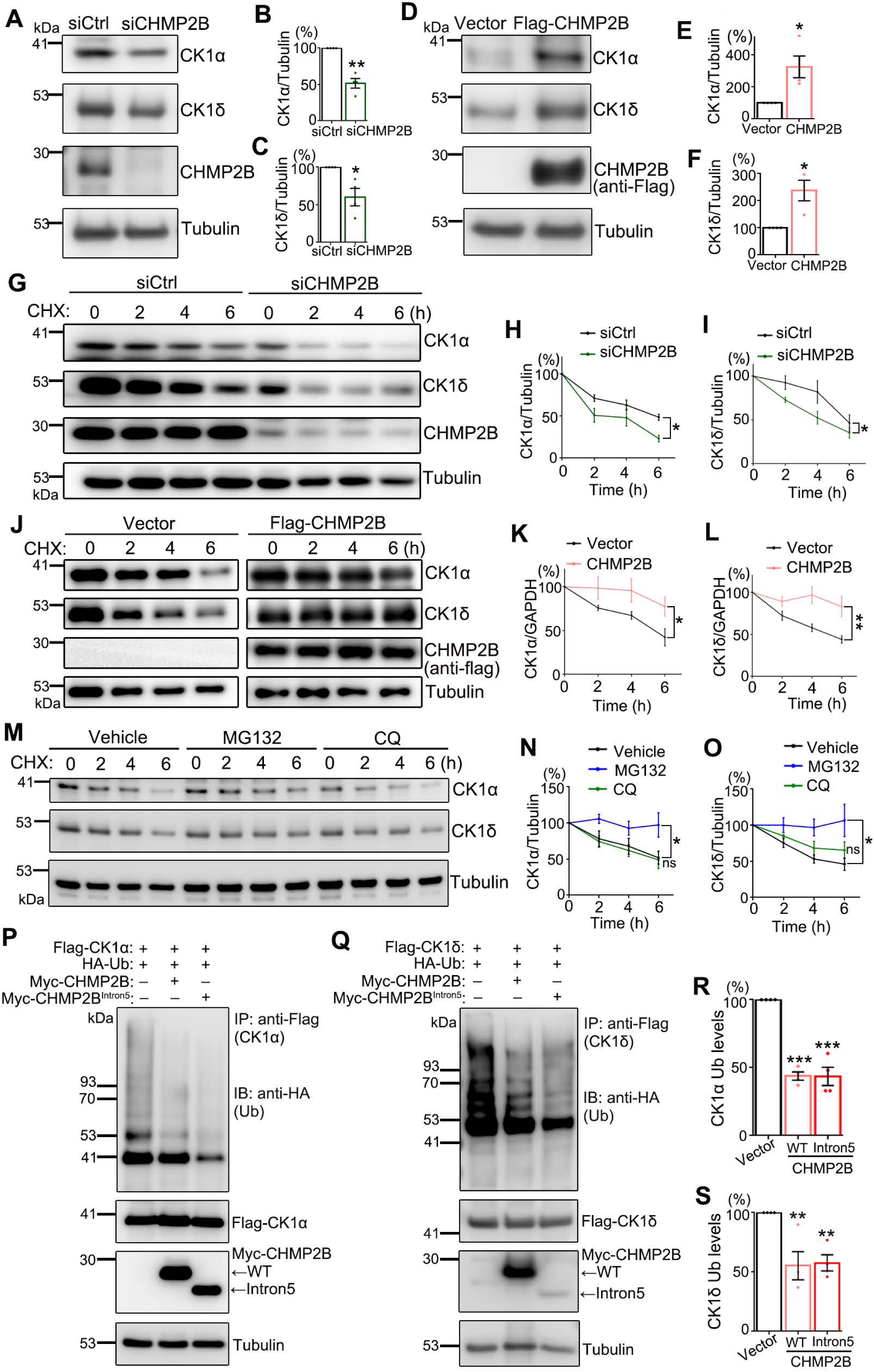
CHMP2B regulates UPS-mediated turnover of CK1α and CK1δ. **(A-F)** KD of CHMP2B (A-C) reduces whereas OE of CHMP2B (D-F) increases the protein abundance of CK1α and CK1δ. **(G-L)** The protein turnover rates of CK1α and CK1δ in 293T cells with CHMP2B KD (G-I) or OE (J-L) are assessed by the pulse chase assay. The time (h) after the cycloheximide (CHX) treatment is indicated. All proteins are normalized to Tubulin and the relative levels at 0 h of each group are set to 100%. **(M-O)** The proteasome inhibitor MG132 (10 μM) but not the autophagy-lysosome blocker CQ (10 μM) significantly suppresses the turnover of CK1α and CK1δ. DMSO is used as a vehicle control. All proteins are normalized to GAPDH and the relative levels at 0 h of each group are set to 100%. **(P-S)** Cells expressing HA-Ubiquitin (HA-Ub) and Flag-tagged CK1α (P, R) or CK1δ (Q, S) are co-transfected with WT or Intron5 CHMP2B as indicated. The Flag-CK1α and Flag-CK1δ proteins are then immunoprecipitated with anti-Flag and the ubiquitination levels are examined with anti-HA by Western blotting. Mean ± SEM, n = 4; \**p* < 0.05, \*\**p* < 0.01, \*\*\**p* < 0.001; ns, not significant. Student’s *t*-test (B-C, E-F), two-way ANOVA (H-I, K-L, N-O), one-way ANOVA (R-S).

Furthermore, we showed that the proteasome inhibitor MG132 but not the autophagy-lysosome inhibitor chloroquine blocked the turnover of CK1α and CK1δ (Figure 5M-5O), indicating that their turnover relied on the UPS-mediated protein degradation but not the autophagy pathway. To determine whether CHMP2B regulated the function of the proteasomal degradation machinery, we conducted an *in vitro* proteasome activity assay. The results showed that neither OE of WT CHMP2B or CHMP2B^intron5^ (Figure S6A) nor KD of CHMP2B (Figure S6B) significantly affected the proteasomal activity of the cells.

Ubiquitination plays an important role in protein degradation, which often serves as a signal for protein disposal via the UPS or autophagy pathway (Khaminets et al., 2016; Kirkin et al., 2009). We performed an *in vitro* proteasome activity assay, which indicated that CHMP2B did not affect the overall proteasomal function in cells (Figure S5). We then tested whether CHMP2B regulated the ubiquitination levels of CK1α or CK1δ. Indeed, we found that both expression of the disease-causal mutation CHMP2B^Intron5^ and increase of WT CHMP2B levels significantly deceased the ubiquitination levels of CK1α and CK1δ (Figure 5P-5S). Together, these data indicate that CHMP2B regulates the ubiquitination and turnover of CK1 via the UPS-dependent protein degradation pathway.

### CHMP2B protein level is upregulated in the brain of aged mice

Aging is a risk factor for neurodegenerative diseases including ALS and FTD, as the onset of the diseases and TDP-43 pathology are age-dependent (Niccoli et al., 2017). In particular, pTDP-43 levels are increased in the brain and spinal cord of WT mice during aging (Liu et al., 2015) and the occurance of individuals with pTDP-43 in the brain increases with age (Riku et al., 2019). Since our cell culture data indicated that increase of CHMP2B impeded CK1 turnover and promoted TDP-43 hyperphosphorylation, we wondered whether increase of CHMP2B levels ever occurred *in vivo* and if aging might have an effect on CHMP2B abundance. Hence, we examined the protein levels of CHMP2B in young adult (2-month old) and aged mice (10-month old). Western blot analyses of the mouse central nervous system demonstrated a striking increase in the protein levels of CHMP2B in the cerebral cortices without the motor cortex (Figure 6A-B), which covered the human brain counterparts of most of the frontal, the entire temporal and parietal lobes. In addition, the protein levels of CHMP2B in the hippocampus of aged mice showed an upward trend with a marginal significance (*p* = 0.056) (Figure 6C-6D). No significant change in the protein levels of CHMP2B was detected in the mouse motor cortex (Figure 6E-6F) or the spinal cord (Figure 6G-6H) during aging. Given the importance of the frontal-temporal lobes and hippocampus in cognitive functions in humans (Burgess et al., 2002; Simons and Spiers, 2003), these data suggest that the age-dependent increase of CHMP2B abundance might be a contributing factor in the development of TDP-43 pathology and the onset of the diseases especially for those associated with dementia.

**Figure 6.**
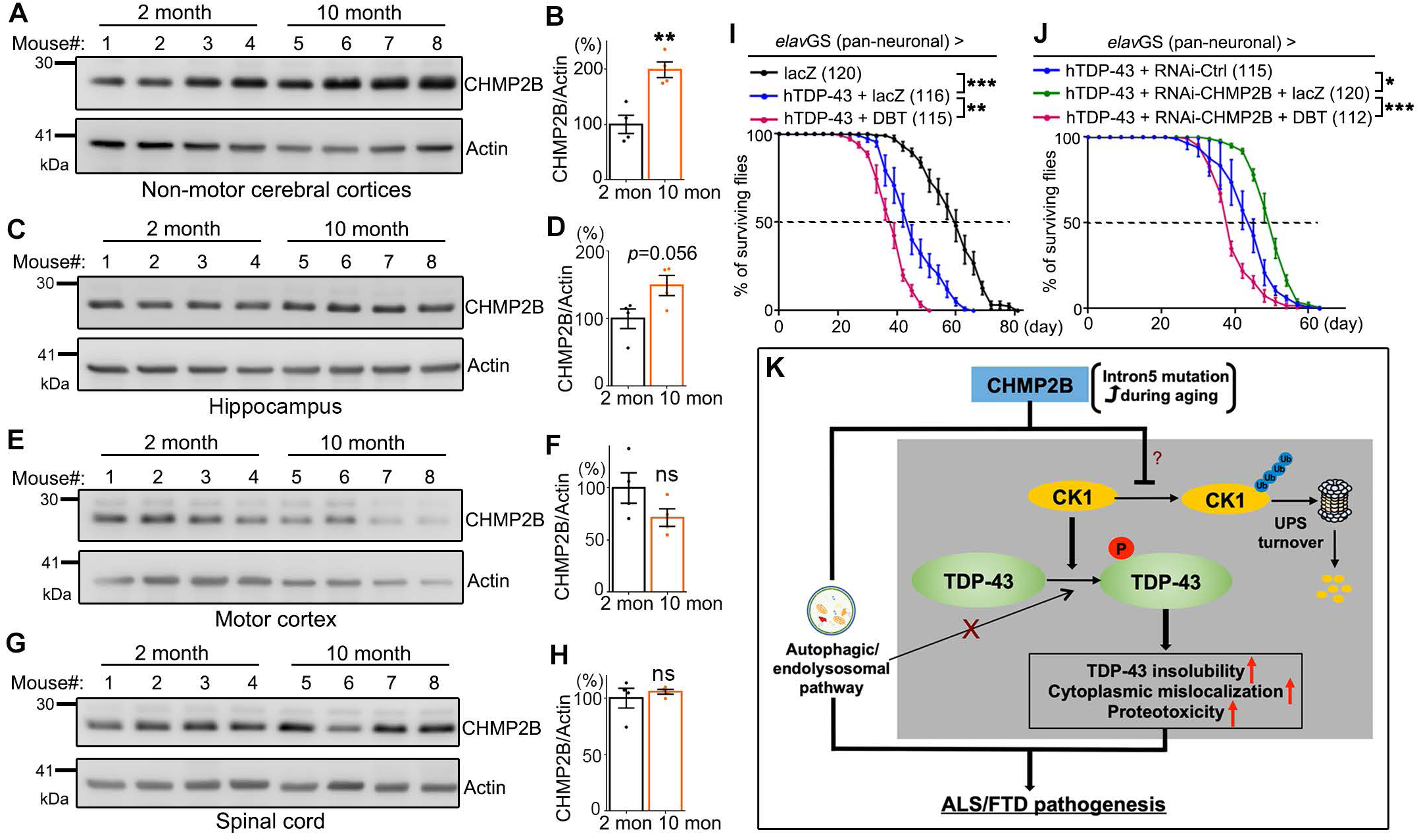
The protein abundance of CHMP2B in various regions of the mouse CNS during aging. The protein levels of CHMP2B in the non-motor cerebral cortices (**A-B**), the hippocampus (**C-D**), the motor cortex (**E-F**) and the spinal cord (**G-H**) of young (2-month) and aged (10-month) mice are analyzed by Western blotting. n = 4 mice per group. (**I-J**) Lifespan assays of the flies with adult-onset, neuronal expression (by *elav*GS) of the transgenes as indicated. The numbers of flies tested in each group are indicated. The UAS-*lacZ* flies are used as a UAS control. Mean ± SEM; \**p* < 0.05, \*\**p* < 0.01, \*\*\**p* < 0.001; ns, not significant; Student’s *t*-test (B, D, F, H) and two-way ANOVA (I-J). (**K**) A schematic model of the “CHMP2B–CK1–TDP-43” pathogenic axis. In addition to the function in the autophagic-endolysosomal pathway, we discover in this study that CHMP2B modulates the ubiquitination levels and the protein turnover of CK1 via the UPS-dependent pathway. In particular, the disease causal mutation CHMP2B^Intron5^ or increased levels of CHMP2B with age may reduce CK1 turnover and increase its protein abundance, which promotes TDP-43 hyperphosphorylation, leading to increased insolubility, cytoplasmic mislocalization and proteotoxicity of TDP-43. Together, CK1-mediated TDP-43 hyperphosphorylation may contribute to the pathogenesis of CHMP2B-related ALS/FTD and other diseases. Future studies are required to examine how CHMP2B regulates the ubiquitination of CK1 and whether the relationship between CHMP2B and pTDP-43 exists in mammalian models *in vivo*, potentially expanding the therapeutic relevance of these findings.

### Increase of CK1 abolishes the mitigating effect of RNAi-CHMP2B on TDP-43 flies

This study was initiated by the identification of RNAi-CHMP2B as a suppressor of TDP-43-induced neurodegeneration in a *Drosophila* screen. Finally, we sought to verify that CK1 mediated the modifying effect of CHMP2B on TDP-43 in *in vivo* settings. Previous studies demonstrated that inhibition of CK1, mainly the δ and ε isoforms, suppressed TDP-43-induced neurotoxicity in fly and mouse models of ALS and FTD (Alquezar et al., 2016; Hicks et al., 2019; Martinez-Gonzalez et al., 2020). Here, we showed that OE of *Doubletime* (*DBT*), the *Drosophila* homologue of CK1δ/ε, in adult fly neurons enhanced the TDP-43 toxicity, as the longevity of TDP-43 flies was further shortened (median life, from 44.3 ± 2.1 d to 39.3 ± 1.4 d) (Figure 6I). More importantly, upregulation of CK1 levels by OE of *dbt* abolished the lifespan-extending effect of RNAi-CHMP2B on TDP-43 flies (median life, from 50.3 ± 0.8 d to 38.5 ± 0.5 d) (Figure 6J). Together with the data from the cell culture-based experiments (Figures 4 and 5), these findings indicated a crucial role of CK1 in mediating the modifying effect of CHMP2B on TDP-43 pathogenesis.

## DISCUSSION

CHMP2B is a major component of the ESCRT-III complex and the function of the ESCRT machinery is required for the biogenesis of multivesicular bodies (MVBs). The ESCRT pathway and the MVBs are not only in charge of sorting and delivering various cellular cargos to vacuoles and lysosomes for degradation, but also regulate a variety of biological processes such as retroviral budding, cytokinetic abscission, and the formation and maintenance of synapses (Henne et al., 2011; Hurley, 2010; Chassefeyre et al., 2015). Rare pathogenic mutations in *CHMP2B* are associated with ALS and FTD (Skibinski et al., 2005; Parkinson et al., 2006; Cox et al., 2010). Previous studies of the *CHMP2B* mutations using human cells as well as fly and mouse models demonstrate a major defect in the autophagic-endolysosomal pathway (Lee et al., 2007; van der Zee et al., 2008; Ghazi-Noori et al., 2012; Clayton et al., 2015; Vernay et al., 2016; Clayton et al., 2018;). Also, misregulation in several major signaling pathways such as Toll (Ahmad et al., 2009), Notch (Cheruiyot et al., 2014), and TGF-β and JNK signaling (West et al., 2015) are assoicated with mutant CHMP2B *in vivo*. In addition, neuronal expression of CHMP2B mutations such as CHMP2B^Intron5^ impairs the maturation of dendrtic spines (Belly et al., 2010), causes inclusion formation and axonal degeneration (Ghazi-Noori et al., 2012), and develops pathological and behavioral features of ALS and FTD (Vernay et al., 2016).

In this study, we identify RNAi-*CHMP2B* as a suppressor of hTDP-43-mediated neurodegeneration in flies, which raised the possibility that its mammalian counterpart is also a potential modifier of TDP-43. Indeed, we show that manipulation of CHMP2B levels by OE or KD of human CHMP2B in 293T cells demonstrates a positive correlation between CHMP2B levels and TDP-43 cytotoxicity. More interestingly, KD of CHMP2B reduces whereas OE of WT or Intron5 CHMP2B increases pTDP-43 levels, pointing to a crucial and previously unknown function of CHMP2B in regulating the phosphorylation states of TDP-43. Furthermore, OE of CHMP2B or CHMP2B^Intron5^ promotes the insolubility and abnormal cytoplasmic localization of TDP-43. The identification of CHMP2B as a modifier of TDP-43 phosphorylation and proteotoxicity led us to examine whether the autophagy-lysosomal pathway is involved. Surprisingly, this regulation is independent of the autophagy-lysosomal pathway, as inhibition of the autophagic or lysosomal function does not alter TDP-43 phosphorylation levels.

CK1 is a known kinase that phosphorylates TDP-43 *in vitro* (Hasegawa et al., 2008; Kametani et al., 2009). We provide evidence that CK1 is a molecular link between CHMP2B and TDP-43, which mediates the modifying effect of CHMP2B on TDP-43 phosphorylation and proteotoxicity. Our findings further show that CHMP2B controls the abundance of CK1 protein by regulating the ubiquitination levels and the UPS-dependent turnover of CK1. It is unclear how exactly CHMP2B regulates the ubiquitination of CK1, which is definitely worth further investigation in the future. Along the line, it is not completely surprising that the ESCRT complex also functions in protein ubiquitination. For example, CHMP5, another ESCRT-III component, has been shown to interact with the deubiquitinase USP8 and regulate the ubiquitination of proteins in immune cells (Adoro et al., 2017; Son et al., 2019).

KD of CK1α or CK1δ can partially rescue the cell viability impaired by CHMP2B OE, while OE of CK1α or CK1δ abolishes the mitigating effect of KD of CHMP2B in 293T cells, which suggest that increased cellular CK1 levels are at least in part responsible for CHMP2B-mediated cytotoxicity. Thus, in addition to autophagy dysfunction and misregulation of the Toll and other cellular signaling pathways, the CK1-mediated hyperphosphorylation of TDP-43 may be another key mechanism contributing to the pathogenesis of CHMP2B-related ALS/FTD. Besides, CK1 is also involved in AD (Ghoshal et al., 1999; Yasojima et al., 2000; Chen et al., 2017) and its colocalization with CHMP2B is found in granulovacuolar degeneration bodies in AD (Funk et al., 2011). Given that TDP-43 pathology is present in up to 57% AD cases (McAleese et al., 2017) and CHMP2B protein levels increase during aging in the mouse brain cortices important for cognitive function (Figure 6A-6B), it is conceivable that the molecular axis of “CHMP2B–CK1– TDP-43” (Figure 6K) may play a broader and more fundamental role in the age-dependent onset and progression of neurodegenerative diseases. And, it will be interesting to determine whether modulation of CK1 activity may serve as a potential therapeutic target for CHMP2B-related diseases and dementia.

## METHODS AND MATERIALS

### *Drosophila* strains

The following strains were obtained from the Bloomington *Drosophila* Stock Center (BDSC): RNAi-*mCherry* (#35785, a control for short hairpin RNAi knockdown), RNAi-*luciferase* (#31603, a control for long hairpin RNAi knockdown), *elav*GS (#43642), RNAi-*CHMP2B* (#28531) and the UAS-*dbt* line (#12121). The RNAi-*CHMP2B* (#38375) strain was obtained from the Tsinghua Fly Center (TFC). For the long hairpin RNAi line of *CHMP2B* (#28531), a copy of UAS-*Dcr2* was co-expressed to boost the knockdown efficiency (Ni et al., 2007). The UAS-*TDP-43* flies were described previously (Sun et al., 2018).

Flies tested in this study were raised on standard cornmeal media and maintained at 25 °C and 60% relative humidity. For adult-onset, neuronal expression of the UAS or RNAi transgenes using the *elav*GS driver (Osterwalder et al., 2001), flies were raised on regular fly food supplemented with 80 µg/ml RU486 (TCI).

### Fly lifespan and climbing assays

For the lifespan experiment, 20 flies per vial and 5-8 vials per group were tested. Flies were transferred to fresh fly food every 3 days and the number of dead flies of each vial was recorded. Flies lost prior to natural death through escape or accidental death were excluded from the final analysis. The median lifespan was calculated as the mean of the medians of each vial belonging to the same group, whereas the “50% survival” shown on the survival curves was derived from compilation of all vials of the group. For the climbing assay, 20 flies were transferred into an empty polystyrene vial and gently tapped down to the bottom of the vial. The number of flies that climbed over a distance of 3 cm within 10 seconds was recorded. The test was repeated three times for each vial and 5-8 vials of each genotype were assessed.

### RNA extraction and real-time quantitative PCR (qPCR)

For qPCR, total RNA was isolated from fly heads or cell culture using TRIzol (Invitrogen) according to the manufacturer’s instruction. After DNase (Promega) treatment to remove genomic DNA, the reverse transcription (RT) reactions were performed using All-in-One cDNA Synthesis SuperMix kit (Bimake). The cDNA was then used for real-time qPCR using 2x SYBR Green qPCR Master Mix (Bimake) with the QuantStudio™ 6 Flex Real-Time PCR system (Life Technologies). The mRNA levels of *actin* or *GAPDH* were used as an internal control to normalize the mRNA levels of genes of interest. The qPCR primers used in this study are listed below:

*dActin* forward: 5’-GAGCGCGGTTACTCTTTCAC-3’

*dActin* reverse: 5’-GCCATCTCCTGCTCAAAGTC-3’

*dCHMP2B* forward: 5’-GAAAGAAACCCACCGTGAAG-3’

*dCHMP2B* reverse: 5’-TCCTCCTCCTCCATTTTCCT-3’

*hβ-actin* forward: 5’-GTTACAGGAAGTCCCTTGCCATCC-3’

*hβ-actin* reverse: 5’-CACCTCCCCTGTGTGGACTTGGG-3’

*hCHMP2B* forward: 5’-AGATGGCTGGAGCAATGTCT-3’

*hCHMP2B* reverse: 5’-CCTTCTGGAAATTCTGCATTG-3’

*hCK1α* forward: 5’-TAATGGGTATTGGGCGTCAC-3’

*hCK1α* reverse: 5’-TGGTATGTGTTGCCTTGTCC-3’

*hCK1δ* forward: 5’-AGCACATCCCCTATCGTGAG-3’

*hCK1δ* reverse: 5’-AGCCCAGAGACTCCAAGTCA-3’

### Plasmids and siRNAs

The pCAG-hTDP-43-HA plasmid was generated as previously described (Sun et al., 2018). To generate pCAG-Flag-CHMP2B, pCAG-Flag-CK1α and pCAG-Flag-CK1δ plasmids, DNA fragments encoding human CHMP2B, CK1α and CK1δ were amplified from pCMV3-Flag-CHMP2B (Sino Biological Inc. #HG14596-NF), or cDNA from 293T cells or SH-SY5Y cells by PCR using primers containing the Flag tag sequence. The PCR products were then sub-cloned into a pCAG vector (Chen et al., 2014) using the XhoI/EcoRI sites. The PCR primers used are listed below:

Flag-CHMP2B-F: 5’--CATCATTTTGGCAAAGAATTCGCCACCATGGATTACAAGGAT--3’

Flag-CHMP2B-R: 5’--GCTCCCCGGGGGTACCTCGAGTTAATCTACTCCTAA--3’

Flag-CK1α-F: 5’--CATCATTTTGGCAAAGAATTCGCCACCATGGATTACAAGGATGACGACGATAAGATGGCGAGTAGCAGC--3’

Flag-CK1α-R: 5’--GCTCCCCGGGGGTACCTCGAGTTAGAAACCTTTCATGTTAC--3’

Flag-CK1δ-F: 5’--CATCATTTTGGCAAAGAATTCGCCACCATGGATTACAAGGATGACGACGATAAGATGGAGCTGAGAGTC--3’

Flag-CK1δ-R: 5’--GCTCCCCGGGGGTACCTCGAGTCATCGGTGCACGAC--3’

The expression construct of CHMP2B^Intron5^ was generated by homologous recombination. Briefly, the DNA fragment of Flag-CHMP2B^Intron5^ was amplified by PCR and inserted into the cloning vector using ClonExpress MultiS One Step Cloning Kit (Vazyme). The construct was then sub-cloned into the pCAG expression vector as above. The PCR primers used in this study: Flag-CHMP2B^Intron5^-F: 5’—CATCATTTTGGCAAAGAATTCGCCACCATGGATTACAAGGAT--3’ Flag-CHMP2B^Intron5^-R: 5’--GCTCCCCGGGGGTACCTCGAGTTACACCTTTCCAGA--3’ The siRNA oligonucleotides to CHMP2B, CK1α and CK1δ were purchased from GenePharma (Shanghai, China), and the sequences of siRNAs are listed below:

si-Ctrl (Negative control): ACGUGACACGUUCGGAGAA

si-CHMP2B#1: UUUAUUACAUCAUCCACGG

si-CHMP2B#2: AGAUGCACAAGUUGUUUGG

si-CHMP2B#3: CAGAUGGUAAGCUUCGAGC

si-CK1α#1: UUCUACUGAUCAUCUGGUC

si-CK1α#2: UAACUGGUUUAAUCCUGAG

si-CK1α#3: UUUCUGCUUUAACAUUGUC

si-CK1δ#1: CAUUUGGUCAGCAAGCAGC

si-CK1δ#2: UUGUUCAAUUCCAAGGUGC

si-CK1δ#3: AUUUCUGUCUCUUGGUGGC

### Cell cultures, transfection

293T cells were cultured in Dulbecco’s Modified Eagle Medium (Sigma-Aldrich, D0819) supplemented with 10% (v/v) Fetal Bovine Serum (BioWest) and GlutaMAX™ (Invitrogen) at 37 °C in 5% CO_2_. Transient transfection of siRNA oligonucleotides or plasmids was performed using Lipofectamine™ RNAiMAX (Invitrogen) in Opti-MEM (Invitrogen), or PolyJet™ In Vitro DNA Transfection Reagent (SignaGen Laboratories) in DMEM, respectively. Cells were harvested 48 h and 72 h after transfection with plasmids and siRNAs, respectively.

### Pharmacological experiments

For the pulse-chase assay, CHX was added into the culture medium to a final concentration of 25 ng/ml. For the proteasomal inhibition, MG132 was added to a final concentration of 10 µM. For the autophagy-lysosome inhibition, chloroquine was added to a final concentration of 10 µM. The cells were treated with the above drugs for indicated durations and then harvested for the subsequent Western blotting or immunocytochemistry analysis.

### Cell viability assay

293T cells were plated in 24-well plates at a density of 200,000 cells/ml, and transfected with siRNAs or plasmids 12-24 h later. Six to eight hours after transfection, the cells were seeded at 10,000 cells/well into 96-well plates at a volume of 100 µl/well. Cell viability was assessed by measuring the reduction of WST-8 [2-(2-methoxy-4-nitrophenyl)-3-(4-nitrophenyl)-5-(2, 4-disulfophenyl)-2H-tetrazolium] into formazan using Cell Counting Kit-8 assay (Dojindo) by adding 10 µl of CCK-8 solution to the cells at specific time points. Thereafter, the cells were incubated for 2 h at 37 °C before measurement of the OD values at 450 nm. Cell viability was quantified according to the manufacturer’s instructions.

### Antibodies

The following antibodies were used for Western blotting, immunoprecipitation and immunofluorescence assays: Rabbit anti-phospho TDP-43 (Ser409/Ser410) (Sigma-Aldrich, SAB4200225), mouse anti-FLAG (Sigma-Aldrich, F3165), mouse anti-Flag (Proteintech, 20543-1-AP), mouse anti-HA (Proteintech, 66006-1), rabbit anti-HA (CST, 3724), rabbit anti-TDP-43 (Proteintech, 10782-2-AP), mouse anti-TDP-43 (Abcam, ab57105), mouse anti-P62/SQSTM1 (Proteintech, 66184-1-Ig), rabbit anti-LC3B (Abcam, ab48394), rabbit anti-CK1α (Proteintech, 55192-1-AP), chicken anti-CK1α (Thermo, PA1-10006), rabbit anti-CK1δ (Proteintech, 14388-1-AP), mouse anti-CK1δ (Thermo, MA5-17243), rabbit anti-GAPDH (Bimake, A5028), mouse anti-GAPDH (Proteintech, 60004-1), rabbit anti-Tubulin (MBL, PM054), mouse anti-actin (Cell Signaling, 3700). HRP conjugated secondary antibodies: anti-mouse (Sigma-Aldrich, A4416), anti-rabbit (Sigma-Aldrich, A9169) and anti-rat (Sigma-Aldrich, A9037). Fluorescent secondary antibodies: anti-Rabbit Cy5 (Life Technologies, A10523), anti-mouse Alexa Fluor^®^ 488 (Life Technologies, A11029) and anti-chicken Alexa^®^ Fluor 633 (Sigma-Aldrich, A-21103).

### Protein extraction and Western blotting

Fly heads or cultured cells were lysed in 2% SDS lysis buffer (100 mM Tris-HCl at pH 6.8, 2% SDS, 40% glycerol, 10% β-mercaptoethanol, 0.04% bromophenol blue) or tissue extraction reagent I (50 mM Tris, pH 7.4, 250 mM NaCl, 5 mM EDTA, 2 mM Na_3_VO_4_, 1 mM NaF, 20 mM Na_4_P_2_O_7_, 0.02% NaN_3_) (Invitrogen) containing protease and phosphatase inhibitor cocktails (Roche, 04693132001). For separation of soluble and insoluble proteins, cells were lysed on ice using RIPA buffer (50 mM Tris pH 8.0, 150 mM NaCl, 1% NP-40, 5 mM EDTA, 0.5% sodium deoxycholate, 0.1% SDS) supplemented with protease and phosphatase inhibitors (Roche). Samples were sonicated and then centrifuged at 13,000 x g for 10 min at 4 °C. The resulting supernatant was used as the soluble fraction and the pellets containing insoluble fractions were dissolved in a 9M urea buffer (9 M urea, 50 mM Tris buffer, pH 8.0) after wash.

All protein samples were boiled at 95 °C for 5 min. Equal mounts of lysates were resolved by electrophoresis using a 10% Bis-Tris SDS-PAGE (Invitrogen) and probed with the primary and secondary antibodies listed above. Detection was performed using the High-sig ECL Western Blotting Substrate (Tanon). Images were captured using an Amersham Imager 600 (GE Healthcare) and the densitometry was measured using ImageQuant TL Software (GE Healthcare) and ImageJ. The contrast and brightness were optimized equally in Adobe Photoshop CS6. Tubulin, GAPDH or Actin was used as a loading control for normalization as indicated in the figures.

### Immunocytochemistry and confocal imaging

293T cells grown on coverslips pre-coated with PLL (Sigma-Aldrich) in a 24-well plate were transfected and treated as described above. The cells were fixed in 4% paraformaldehyde in PBS for 15 min at room temperature, permeabilized with 0.5% Triton X-100 (Sigma-Aldrich) in PBS for 15 min, and blocked with 3% goat serum in PBST (0.1% Triton X-100 in PBS) for 1 h at room temperature. The above primary and secondary antibodies in the blocking buffer were then incubated at 4 °C overnight or at room temperature for 1 h. After 3 washes with PBST, cells were mounted on glass slides using VECTASHIELD Antifade Mounting Medium with DAPI (Vector Laboratories).

Fluorescent images were taken with Leica TCS SP8 confocal microscopy system using a 63X oil objective (NA=1.4). Images were assembled into figures using Adobe Photoshop CS6.

### Immunoprecipitation

293T cells were lysed in an IP buffer (50mM Tris-Hcl PH 7.4, 150mM Nacl, 1% NP-40, 1mM EDTA, 5% glycerol) containing protease inhibitor cocktails and nethylmaleimide (inhibit deubiquitination). Following centrifugation at 15000 x g for 15 min at 4 °C, the supernatants were collected in new vials, and incubated with mouse anti-Flag beads on a rotary shaker at 4 °C overnight. The beads were then collected and eluted in the 2X SDS buffer for the subsequent Western blotting assay.

### *In vitro* proteasome activity assay

Proteasomal activity was measured using a Proteasome Activity Assay kit (Abcam, ab107921). 293T cells were transfected with indicated plasmids for 48 h and siRNA for 72 h, followed by trypsin digestion and quantification. The cells were subsequently lysed in 90 μl lysis buffer (0.5% NP40 in PBS), and the lysates were centrifuged at 16000 x g for 15 min at 4 °C and then the supernatants were collected in new vials. The proteasomal activity of the supernatants was determined by assaying the cleavage of a fluorogenic peptide substrate Suc-LLVY-AMC according to the manufacture’s instruction. The substrate peptides were incubated with the cell lysates at 37 °C for 1 h, and the fluorescence intensity was measured at the end of the assay using a microplate reader (BioTek, Ex/Em = 350/440 nm).

### Statistical analysis

Statistical significance in this study is determined by one-way analysis of variance (ANOVA) with Tukey’s HSD post-hoc test, two-way ANOVA with Bonferroni’s post-hoc test, or unpaired, two-tailed Student’s *t*-test with equal variance at \**p* < 0.05, \*\**p* < 0.01, and \*\*\**p* < 0.001 as indicated. Error bars represent the standard error of the mean (SEM).

## DECLARATIONS

## Acknowledements

We thank the BDSC for providing the fly strains, Z. Zhang for the cloning vectors and plasmids, S. Zhang for technical supports, members of the Fang lab for helpful discussion, and J. Yuan for comments and critical reading of the manuscript.

## Author’s contributions

X.S., A.L. and Y.F. conceived the research; X.S., X.D., Y.C. and Y.F. designed the experiments; X.S., X.D., R.H., Y.D., J.C., J.N. and Q.W. performed the experiments; X.S., X.D., R.H. and Y.D., contributed important new reagents; X.S., X.D., K.Z. and Y.F. analyzed the data and interpreted the results; X.S., X.D. and Y.F. prepared the figures; and X.S., X.D., A.L. and Y.F. wrote the paper. All authors read and approved the final manuscript.

## SUPPORTING INFORMATION

**Figure S1.**
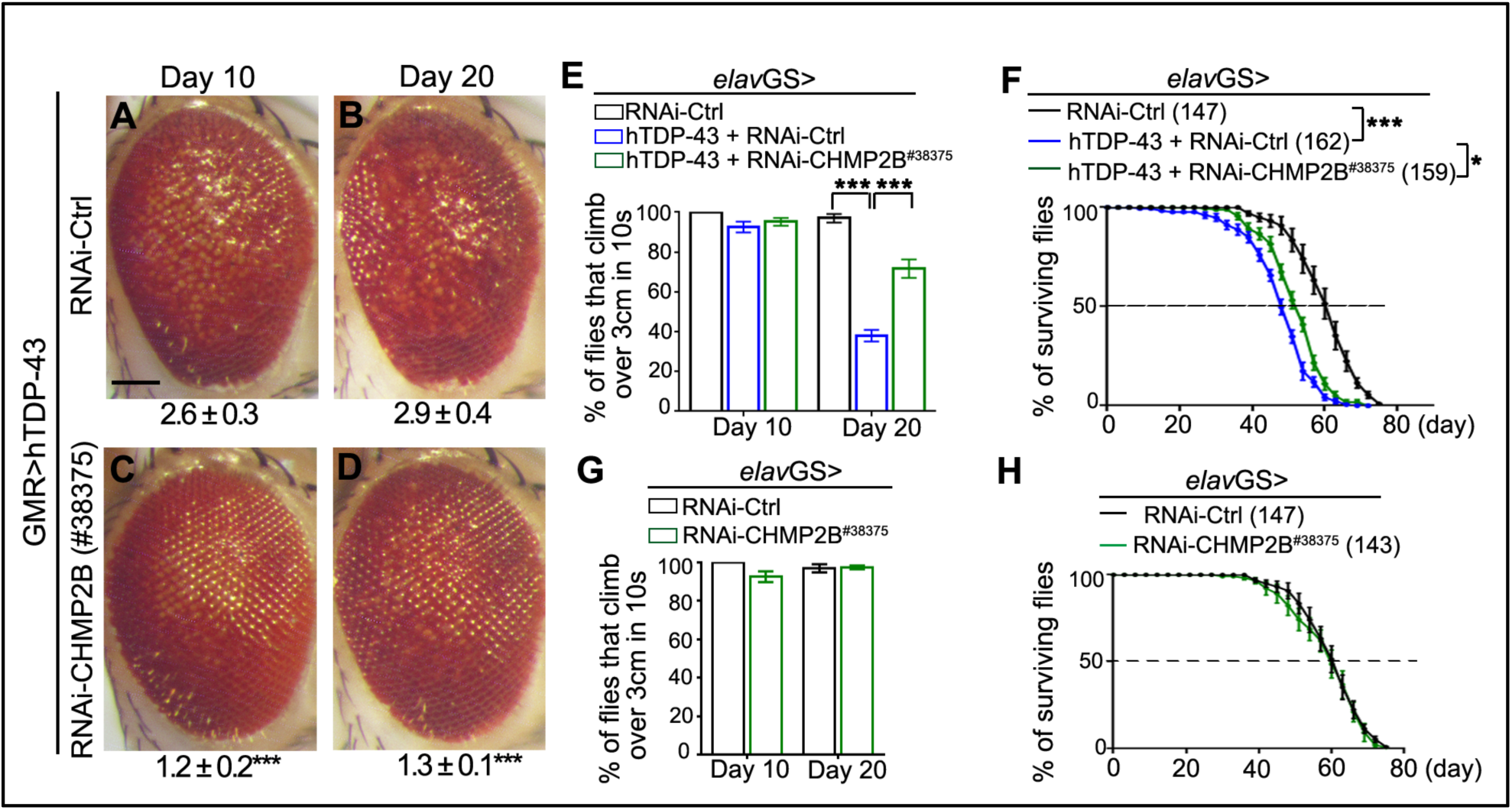
Another independent transgenic RNAi-CHMP2B fly strain (#38375) also exhibits suppression of TDP-43-mediated neurotoxicity. **(A-D)** Representative z-stack images of the eyes of hTDP-43 flies expressing RNAi-Ctrl or RNAi-CHMP2B (#38375) (by GMR-Gal4) at indicated ages. The average degeneration score (mean ± SEM) and the statistical significance compared to the RNAi control line (RNAi-*mCherry*) are indicated at the bottom of each group. **(E-F)** The climbing (E) and lifespan (F) assays of the hTDP-43 flies with RNAi-CHMP2B expressed in adult neurons (by *elav*GS). **(G-H)** The climbing (G) and lifespan (H) assays of the RNAi-CHMP2B flies. No significant difference is detected compared to the RNAi-Ctrl flies. n = 10 eyes/group (A-D), ∼20 flies/vial x 10 vials/group in (E, G), and the numbers of flies tested in each group are indicated in (F, H). Mean ± SEM. \*\**p* < 0.01 and \*\*\**p* < 0.001; Student’s *t*-test (A-E, G) and two-way ANOVA (F, H). Scale bar: 100 μm

**Figure S2.**
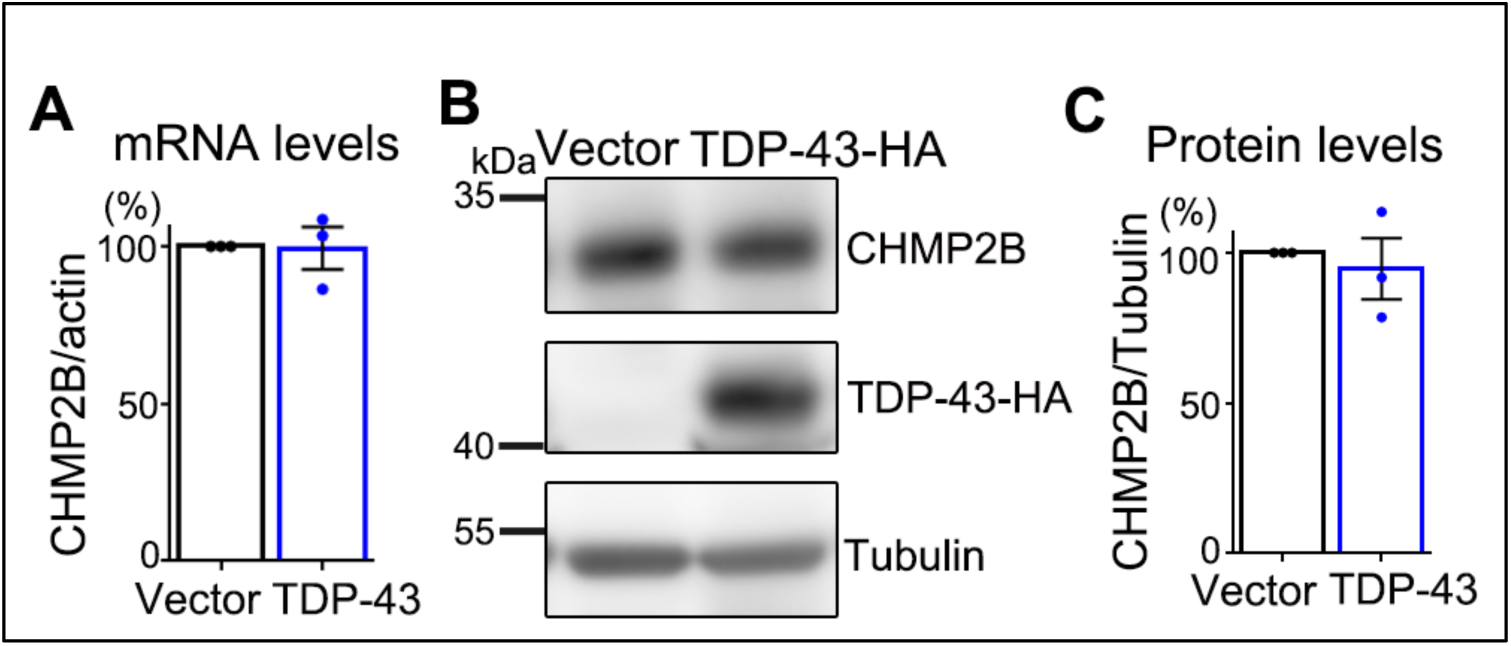
OE of TDP-43 does not alter CHMP2B levels in mammalian cells. **(A)** The mRNA levels of *CHMP2B* in 293T cells transfected with the empty vector or TDP-43-HA are assessed by qPCR. The mRNA levels of *CHMP2B* are normalized to *actin* and shown as average percentages to the vector control group. **(B-C)** The protein levels of CHMP2B in 293T cells transfected with the empty vector or TDP-43-HA are examined by Western blotting. The protein levels of CHMP2B are normalized to Tubulin and shown as average percentages to the control group (Vector). Means ± SEM; n = 3; Student’s *t*-test.

**Figure S3.**
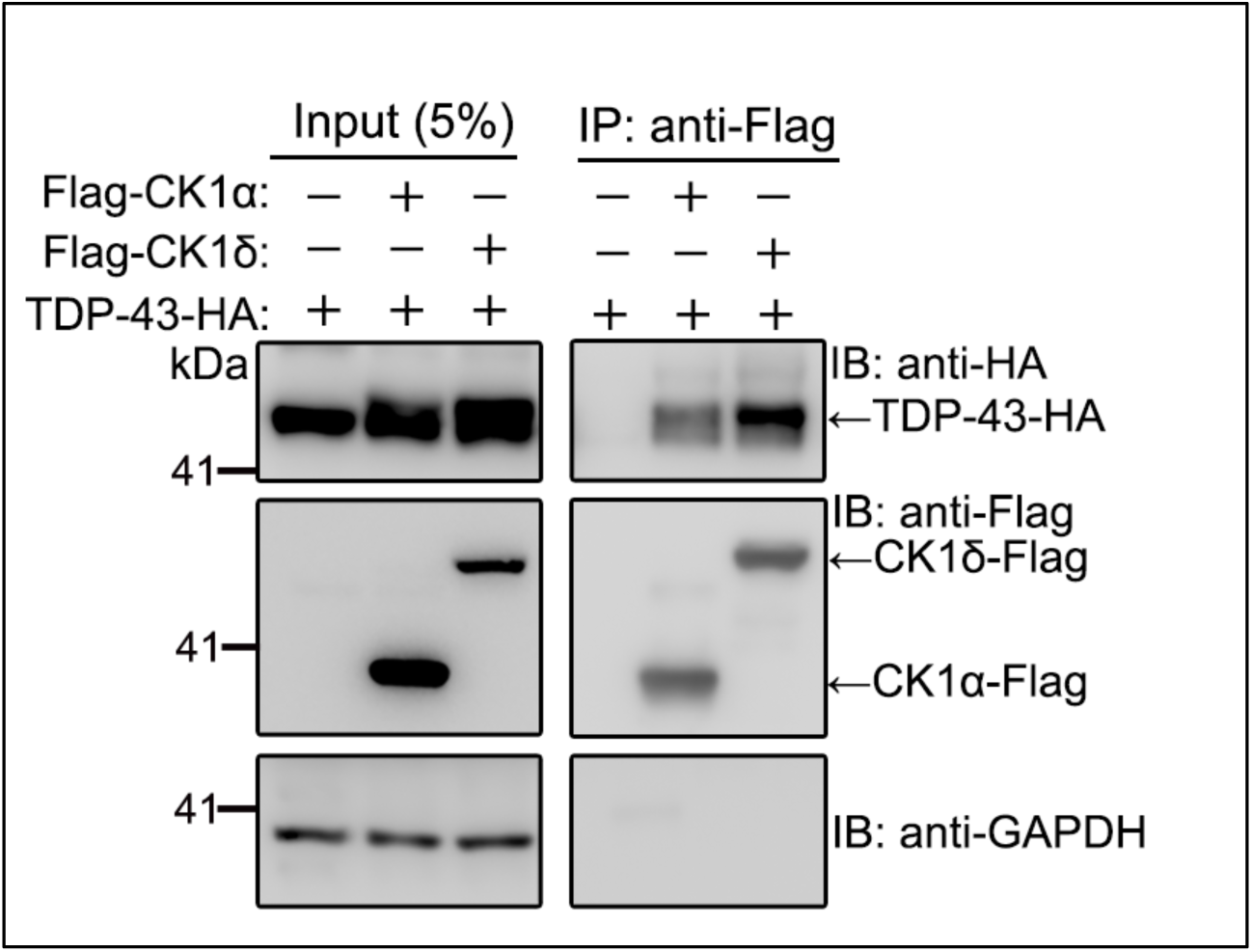
CK1 can interact with TDP-43 in 293T cells. 293T cells expressing TDP-43-HA are co-transfected with the empty vector, Flag-tagged CK1α or CK1δ and the cell lysates are immunoprecipitated with anti-Flag and examined by Western blotting with the antibodies as indicated. The co-immunoprecipitation experiments are independently repeated for 3 times

**Figure S4.**
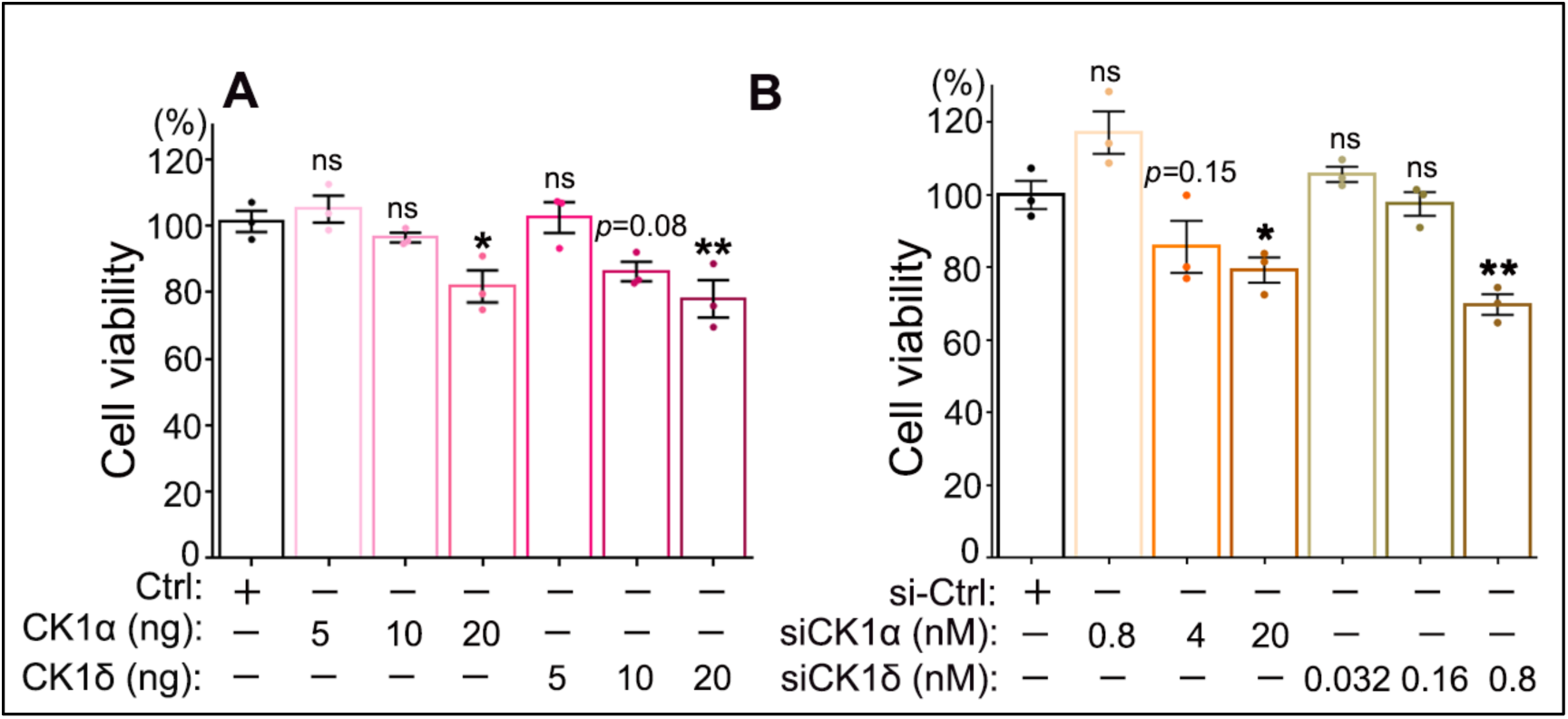
Moderate OE or KD of CK1 does not cause significant cytotoxicity. **(A)** The viability of 293T cells transfected with Flag-CK1α or Flag-CK1δ at indicated concentrations is examined by the CCK-8 assay. Cells transfected with the empty vector are used as a control (Ctrl). **(B)** The viability of 293T cells treated with siCK1α or siCK1δ at indicated concentrations are assessed by the CCK-8 assay. The scrambled siRNA is used as a control (siCtrl). Means ± SEM; n = 3; \**p* < 0.05, \*\**p* < 0.01; ns: not significant; one-way ANOVA.

**Figure S5.**
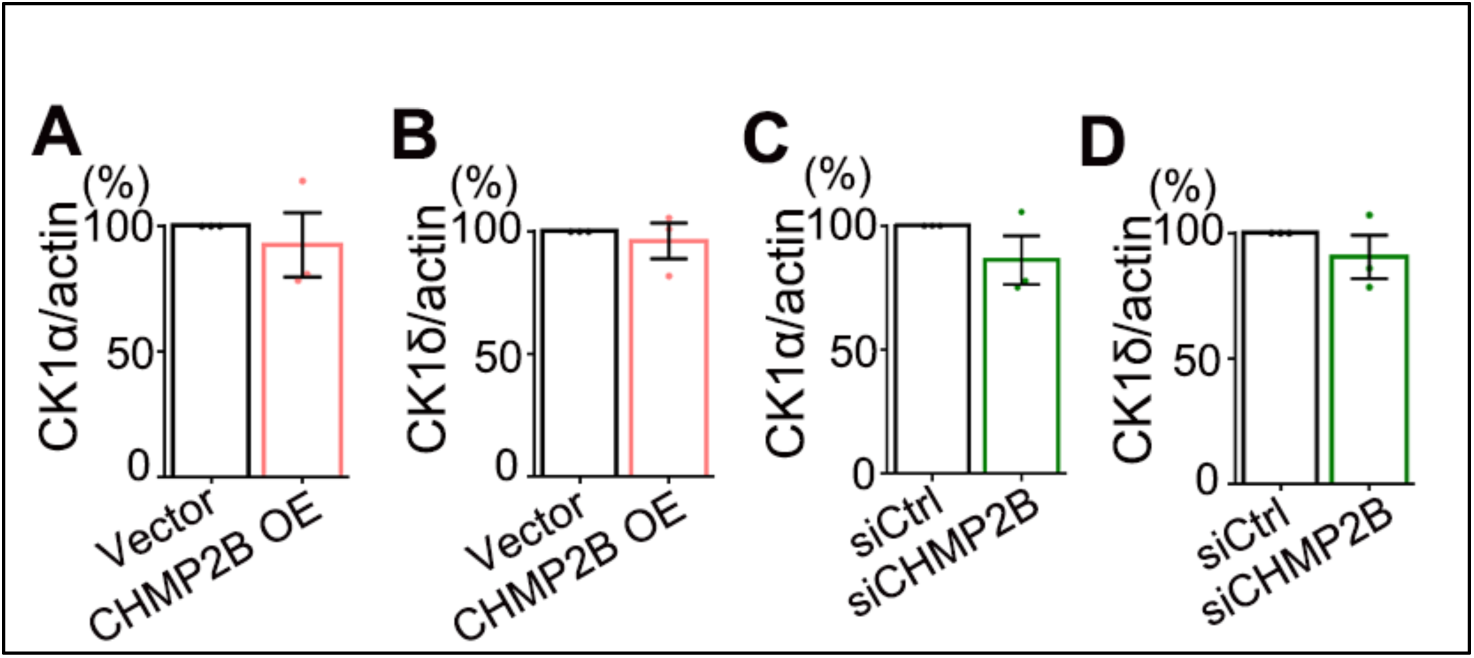
The impact of OE or KD of CHMP2B on the mRNA levels of CK1α and CK1δ. **(A-D)** qPCR analysis of the mRNA levels of CK1α and CK1δ in 293T cells transfected with Flag-CHMP2B (A-B) or siCHMP2B (C-D). All mRNA levels are normalized to *actin* and shown as average percentages to the vector or siRNA control group. Means ± SEM; n = 3; Student’s *t*-test.

**Figure S6.**
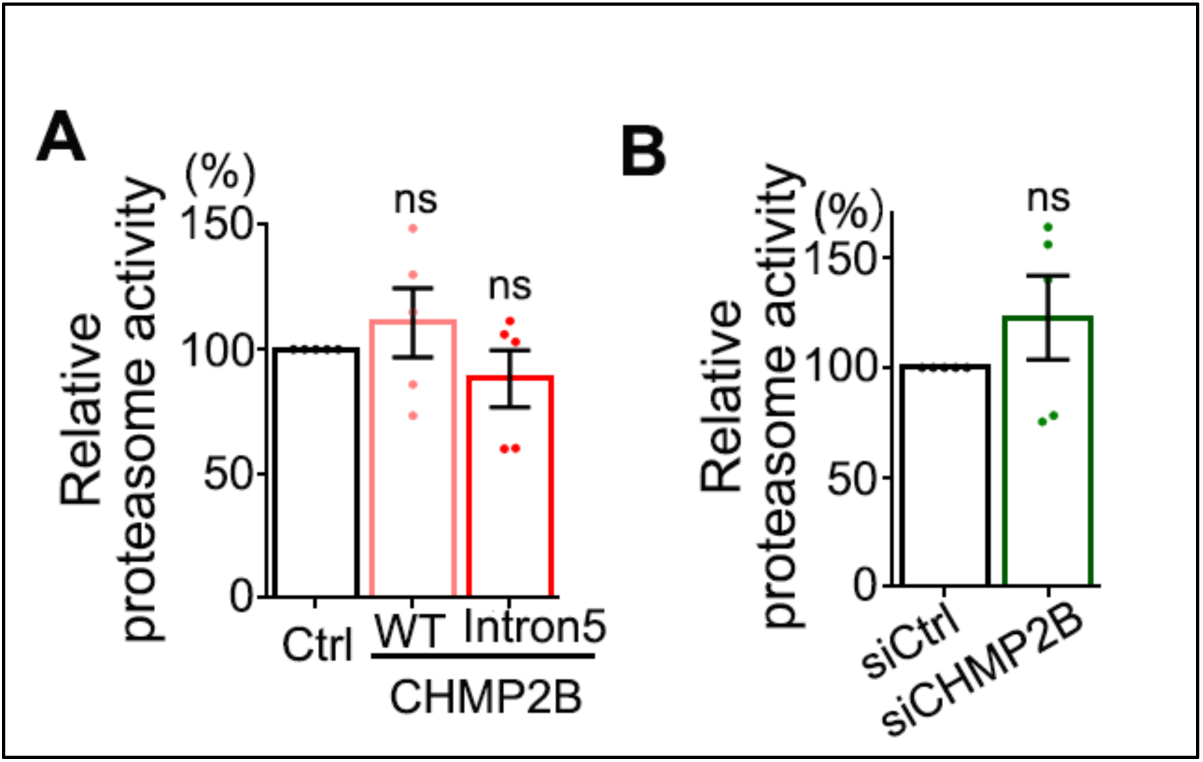
OE or KD of CHMP2B does not significantly alter the cellular proteasome activity. **(A-B)** Relative proteasome acitivity of 293T cells transfected with the empty vector (Ctrl), WT or Intron5 CHMP2B (A) or treated with scrambled siRNA (siCtrl) or siCHMP2B (B). The proteasome activity is determined using an *in vitro* fluorogenic peptide cleavage assay. The relative proteolytic activities are shown as average percentages to the total fluorescence intensity of the control group at the end of the assay (set to 100%). Means ± SEM; n = 5; ns: not significant; one-way ANOVA in (A) and Student’s *t*-test in (B).

